# Multi-Modal Protein Representation Learning with CLASP

**DOI:** 10.1101/2025.08.10.669533

**Authors:** Nicolas Bolouri, Joseph Szymborski, Amin Emad

## Abstract

Effectively integrating data modalities pertaining to proteins’ amino acid sequences, three-dimensional structures, and curated text-based descriptions of their biochemical and functional properties can lead to informative representations capturing different views of proteins. Here, we introduce CLASP, a unified tri-modal framework that combines the strengths of geometric deep learning, natural large language models (LLMs), protein language models (pLMs), and contrastive learning to learn informative protein representations based on their structure, amino acid sequence, and text-based biochemical and functional descriptions. We show that CLASP enables accurate zero-shot classification and retrieval tasks, such as matching a protein structure to its sequence or description, outperforming state-of-the-art baselines. CLASP embeddings also exhibit superior clustering by protein family, and ablation studies confirm that all three modalities contribute synergistically to performance. Our results highlight the power of integrating structural, sequential, and textual signals in a single model, establishing CLASP as a general-purpose embedding framework for protein understanding.

## 1 Introduction

Advances in deep learning have led to powerful protein language models (pLMs), which learn rich representations from large unlabeled sequence databases. Similar to their natural large language model (LLM) counterparts, pLMs are trained on millions of amino acid sequences based on masked language modeling or causal language modeling (e.g., ESM3 [1], ProtT5 [2], SqueezeProt [3]). However, proteins are more than linear amino acid strings; their three-dimensional structures and biochemical contexts are crucial for their function [4]. In fact, protein structures tend to be more conserved than sequences, with structural comparisons revealing remote homology and functional relationships that are invisible to sequence-based methods [5]. The recent proliferation of experimentally determined and computationally predicted protein structures presents a unique opportunity to enhance protein representations with structural information [6]. High-resolution experimental techniques such as X-ray crystallography, nuclear magnetic resonance (NMR) spectroscopy, and cryo-electron microscopy supply atomic-level structural information and coordinates of proteins, which are ultimately captured as Protein Data Bank (PDB) files [7].

In addition to important signals present in proteins’ amino acid sequence and structure, their biochemical properties and functions are often described and curated in natural language, through research articles, biological databases, and annotation resources like UniProt [8]. These textual descriptions capture high-level semantic knowledge, including cellular roles, disease associations, and molecular activities (e.g., ligand-binding properties), which may not be directly inferable from structure or sequence. Integrating such unstructured text into protein representations offers a promising route to bridge the gap between low-level molecular data and the rich, human-interpretable understanding of protein function [9].

To enrich protein representations, various methods have begun combining structure and sequence modalities. Graph-based deep learning has proven effective for modeling protein 3D structures. For example, Progres [6] embeds protein structures as graphs via a simple graph neural network (GNN) and contrastive learning, yielding fixed-length vectors that enable fast structure searches at scale [6]. Such embeddings can be compared in constant time, dramatically speeding up remote fold detection across massive structure databases [6]. Unsupervised pretraining on protein tertiary structures has also been explored; for instance, Chatzianastasis et al. proposed a self-supervised scheme where 3D graph neural networks are trained to predict geometric relationships between protein subgraphs and the global structure, leading to improved downstream classification performance [10]. Recent work such as DeepUrfold employs latent-space representations derived from hybrid generative and convolutional architectures to uncover distant relationships across protein fold space, stressing the importance of representations that capture sequence–structure–function relationships beyond purely geometric similarity [11]. E(n)-invariant GNN (EGNN) architectures have been introduced to better capture molecular geometry [12].

These models enforce rotational and translational symmetry in 3D space, ensuring that learned representations depend only on the intrinsic shape and not on an arbitrary orientation [12]. Such E(n)-invariant networks have demonstrated state-of-the-art performance in molecular property prediction, enabling more robust protein structure embeddings [12]. Overall, these advances illustrate the value of integrating structural information into protein representations; however, purely geometric approaches often overlook other rich data modalities, such as free-text functional annotations.

In parallel, there is a growing interest in multi-modal models that align protein sequences with textual descriptions or annotations [9]. The motivation is that natural language can encode functional and contextual information that may not be readily inferable from sequence alone. Early efforts, such as OntoProtein [13], augmented sequence embeddings with knowledge graph information from biological databases, effectively injecting curated textual knowledge into learned protein representations [13]. More recently, several groups have drawn inspiration from the CLIP (Contrastive Language-Image Pre-training) [14] paradigm, which has achieved remarkable image-text alignment via contrastive learning, to pair proteins with their corresponding descriptions (e.g., ProteinCLIP [15], ProtCLIP [9]). On the other hand, Xu et al. used biomedical ontology terms and descriptions to guide protein language models (ProtST) [16], while Liu et al. introduced a multi-modal framework (ProteinDT) that aligns sequence embeddings with free-text functional descriptions for protein design applications [17]. These approaches demonstrate that language-supervised protein models can be transferred to a variety of tasks, ranging from property prediction to semantic similarity inference [9]. However, most prior protein biotext models are bimodal. They align sequences with text only, using objectives adapted from image-text CLIP [9], but do not explicitly incorporate 3D structural information during pretraining despite the known importance of structure for function. This highlights a need to build tri-modal representation learning strategies that integrate sequence, structure, and textual descriptions, allowing models to link the physical determinants of protein function with the semantic, often higher-level biological contexts described in natural language. Such integration can facilitate more interpretable representations and support tasks that span both mechanistic and functional domains, such as retrieving disease-relevant proteins from structural similarity or predicting functional effects grounded in both geometry and literature [4].

Here, we introduce CLASP (Contrastive Language–Amino acid Sequence–Structure Pretraining), a unified tri-modal framework for protein representation learning. CLASP jointly embeds a protein’s structure (captured through PDB representation), amino acid sequence, and natural language description into a shared vector space by leveraging a contrastive learning objective inspired by CLIP Goes 3D (CG3D) [18]. Empirically, we demonstrate that CLASP’s joint representations enable accurate zero-shot classification and retrieval across modalities – for example, accurately matching a sequence with its structure or description – outperforming other models limited to one or two modalities. Moreover, CLASP’s structure and projected sequence embeddings exhibit clear clustering by protein family and functional class, indicating that the model learns biologically meaningful structure-function relationships. Finally, ablation studies demonstrate that removing any single modality during training leads to a marked drop in performance, underscoring the complementary value of sequence, structure, and text in capturing protein semantics. Together, these findings suggest that CLASP offers a biologically grounded, modality-agnostic embedding space that unifies structural, sequential, and functional views of proteins, laying the groundwork for more interpretable, general-purpose models in protein science.

## 2 Results

### 2.1 CLASP learns protein representations using multi-modal contrastive learning, geometric deep learning and large language models

CLASP (Contrastive Language-Amino acid sequence-Structure Pretraining) is a deep learning model to learn protein representations using multi-modal data. It combines the strength of natural large language models (LLMs), protein language models (pLMs), geometric deep learning, and contrastive learning to learn informative protein representations based on their structure, amino acid sequence, and description (Figure 1). CLASP serves as a unified framework for learning multi-modal protein representations, achieving representative embeddings by jointly optimizing all components through contrastive learning.

**Figure 1.**
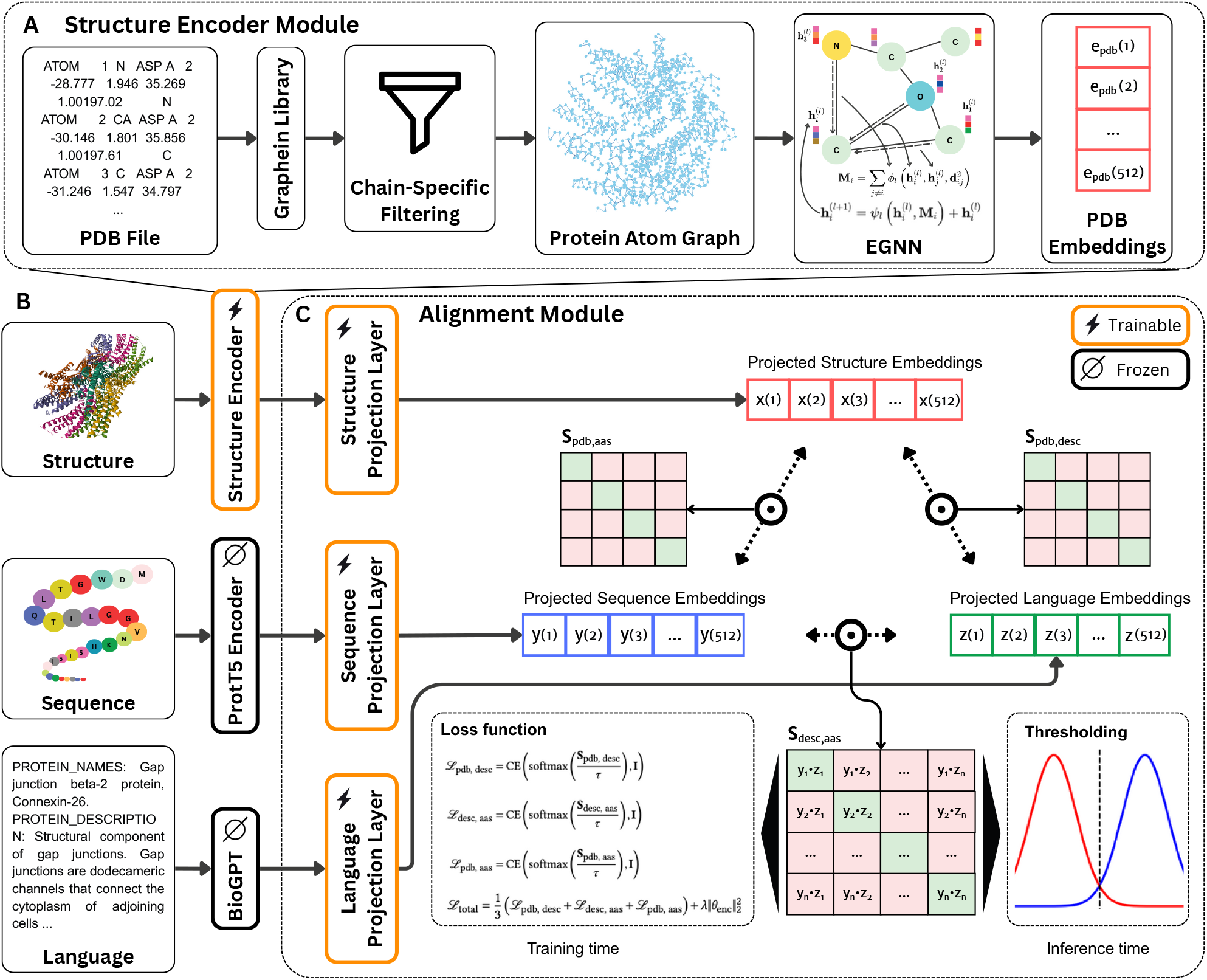
Architecture and pipeline of CLASP. **(A)** The structure encoder module converts protein structures from PDB files into atom-level graphs using the Graphein library, applying chain-specific filtering. Each node is annotated with Meiler descriptors and 3D coordinates, and edges represent spatial proximity. These graphs are processed by an E(3)-invariant Graph Neural Network (EGNN), producing 512-dimensional structure embeddings that preserve geometric and biochemical features. **(B)** Each protein is represented across three modalities: structure (PDB), amino acid sequence, and natural language description. Sequence embeddings are obtained from ProtT5 (frozen) and description embeddings from BioGPT (frozen). These embeddings, along with those from the EGNN, form the inputs to the alignment module. **(C)** The alignment module projects the inputs from each modality into a shared embedding space via learnable linear layers. Pairwise dot-product similarity matrices are computed between all modality pairs and a three-dimensional contrastive learning approach is used to align them together. The entire framework is trained end-to-end using a tri-modal contrastive loss. At inference, thresholding on pairwise similarity matrices enables classification and retrieval tasks.

CLASP’s architecture is comprised of two main components. The first component is a structure encoder module that utilizes geometric deep learning to generate low-dimensional embeddings from the proteins’ PDB files [7] (Figure 1A). This module first converts the PDB file of a protein into a graph representation using Graphein [19]. In this graph, nodes represent the atoms (annotated with biochemical features) and edges describe their spatial relationships. Edge weights in this graph represent Euclidean distances. These graphs are then processed by an E(3)-invariant graph neural network (EGNN) [12], which generates low-dimensional structure embeddings that are invariant to rotations or translations and preserve the protein’s geometric and structural properties. These properties arise from the architecture’s use of distance-based message passing and coordinate updates that are explicitly designed to be invariant to rotations and translations [12]. Such invariance is crucial for molecular structures, where both biochemical and geometric contexts significantly influence function. The EGNN architecture combines multiple message-passing layers to capture both local and global patterns in protein graphs.

The second component is an alignment module that utilizes three-dimensional contrastive learning to symmetrically align structure embeddings with precomputed amino acid sequence and natural language description embeddings in a shared embedding space (Figure 1B). We used precomputed 1024-dimensional ProtT5 [2] amino acid sequence embeddings from UniProt [8]. ProtT5 is a pLM that models protein sequences using a transformer architecture trained on large-scale sequence databases in a masked language modeling framework, enabling it to capture contextual and evolutionary information from amino acid sequences. Protein descriptions were also compiled from UniProt [8], which consisted of possible names and functional descriptions of the proteins. These descriptions were then processed by BioGPT [20], a biology-tuned LLM, generating 1024-dimensional natural language embeddings.

The structure embeddings, along with the precomputed amino acid sequence and language description embeddings go through modality-specific linear projection layers of the same output dimensions (here 512), which map them into a shared embedding space. The outputs of each projection layer are then aligned using an objective function that includes four terms: three symmetric cross-entropy contrastive terms each tasked with aligning embeddings from pairs of modalities and a fourth regularization term. Using this loss function, CLASP learns a unified embedding space in which structure–sequence–description triplets corresponding to the same protein are brought into close proximity, while embeddings from non-matching triplets are explicitly separated, thereby capturing cross-modal correspondences. The CLASP model in its entirety was trained end-to-end, with the contrastive training loss being backpropagated all the way to the structure encoder. The details of this model are discussed in Methods.

### 2.2 CLASP enables accurate alignment of sequence and structure

We first evaluated CLASP’s ability to align amino acid sequences and their corresponding 3D protein structures using a zero-shot classification. Specifically, given a candidate sequence-structure pair, similarity scores were computed via the dot product between their projected embeddings in the shared space to determine whether they belong to the same protein or not.

We randomly split the data into training, validation, and test sets such that proteins in each set would not share more than 30% identity with proteins of other sets (Methods). The model was trained on the training set and the similarity scores were evaluated using the held-out validation set to obtain an optimal decision threshold based on the F1 score. Performance metrics were then computed on the held-out test set using this threshold (repeated three times across distinct randomly formed training/validation/test sets)..

We compared the performance of CLASP on this task with eight alternative methods. Using embeddings provided by two structure embedding models, Progres [6] and COLLAPSE [21], we trained standard CLIP models, Random Forest (RF) classifiers, and logistic regression (LR) models on each embedding set (a total of six different methods). In addition, we evaluated ProstT5 [22], a bilingual protein language model that extends ProtT5 by learning a joint representation of amino acid sequences and discrete structural 3Di sequences, enabling translation between sequence and structure. We used its both translation directions, sequence→structure (AAS→3Di) and structure→sequence (3Di→AAS). Candidate pairs were scored according to the Smith-Waterman alignment [23] between the generated and ground-truth sequences. Decision thresholds were selected on the validation set in the same manner as for CLASP. The same training/validation/test splits were used for all methods to ensure a fair comparison.

As reported in Table 1 and Figure 2A, CLASP substantially outperformed all tested models across multiple evaluation criteria. These results reflect the advantage of the CLASP structural encoder over other encoders, coupled with contrastive tri-modal training to ensure modality consistency. While Progres-CLIP and COLLAPSE-CLIP also adopt contrastive learning objectives, they lag behind CLASP by margins of approximately 5%–7% in Area Under the Receiver Operating Characteristic Curve (AUROC) and Area Under the Precision–Recall Curve (AUPRC), underscoring the contribution of geometry-aware encoding and the benefits of tri-modal alignment in achieving robust structure-sequence alignments. Across both tasks, ProstT5 lags behind CLASP by sizable margins, with MCC gaps of 0.161 for Structure→Sequence and 0.230 for Sequence→Structure. Traditional classifier baselines, such as LR and RF trained on the same embeddings, also underperformed relative to their CLIP-based counterparts, highlighting the strength of contrastive approaches in this setting.

**Table 1.** Classification performance of all models across sequence-structure (AAS-PDB) and description-structure (DESC-PDB) alignment tasks.

| Task | Model | Accuracy | F1 Score | AUROC | AUPRC | MCC |
| --- | --- | --- | --- | --- | --- | --- |
| AAS-PDB | CLASP | **0.920 ± 0.003** | **0.920 ± 0.003** | **0.976 ± 0.002** | **0.977 ± 0.002** | **0.841 ± 0.006** |
|  | Progres - CLIP | 0.843 ± 0.003 | 0.850 ± 0.002 | 0.919 ± 0.002 | 0.914 ± 0.003 | 0.689 ± 0.006 |
|  | ProstT5 (3Di → AAS) | 0.831 ± 0.006 | 0.809 ± 0.007 | 0.855 ± 0.003 | 0.903 ± 0.003 | 0.680 ± 0.012 |
|  | ProstT5 (AAS → 3Di) | 0.800 ± 0.006 | 0.780 ± 0.007 | 0.831 ± 0.003 | 0.882 ± 0.002 | 0.611 ± 0.016 |
|  | COLLAPSE - CLIP | 0.784 ± 0.006 | 0.799 ± 0.001 | 0.869 ± 0.002 | 0.865 ± 0.006 | 0.574 ± 0.009 |
|  | Progres - RF | 0.756 ± 0.001 | 0.734 ± 0.001 | 0.846 ± 0.000 | 0.853 ± 0.001 | 0.519 ± 0.002 |
|  | COLLAPSE - RF | 0.679 ± 0.001 | 0.680 ± 0.004 | 0.746 ± 0.001 | 0.731 ± 0.010 | 0.359 ± 0.003 |
|  | COLLAPSE - LR | 0.583 ± 0.009 | 0.596 ± 0.014 | 0.617 ± 0.002 | 0.587 ± 0.005 | 0.167 ± 0.018 |
|  | Progres - LR | 0.578 ± 0.004 | 0.583 ± 0.008 | 0.614 ± 0.005 | 0.599 ± 0.004 | 0.157 ± 0.007 |
| DESC-PDB | CLASP | **0.770 ± 0.011** | **0.791 ± 0.005** | **0.858 ± 0.007** | **0.846 ± 0.007** | **0.552 ± 0.018** |
|  | Progres - RF | 0.694 ± 0.005 | 0.649 ± 0.006 | 0.777 ± 0.008 | 0.792 ± 0.007 | 0.401 ± 0.011 |
|  | Progres - CLIP | 0.687 ± 0.010 | 0.729 ± 0.001 | 0.778 ± 0.005 | 0.771 ± 0.010 | 0.394 ± 0.009 |
|  | COLLAPSE - CLIP | 0.671 ± 0.013 | 0.716 ± 0.005 | 0.756 ± 0.003 | 0.746 ± 0.003 | 0.361 ± 0.016 |
|  | COLLAPSE - RF | 0.606 ± 0.019 | 0.620 ± 0.021 | 0.654 ± 0.015 | 0.642 ± 0.012 | 0.213 ± 0.038 |
|  | COLLAPSE - LR | 0.557 ± 0.010 | 0.563 ± 0.008 | 0.585 ± 0.012 | 0.551 ± 0.009 | 0.115 ± 0.020 |
|  | Progres - LR | 0.551 ± 0.011 | 0.550 ± 0.008 | 0.573 ± 0.012 | 0.565 ± 0.010 | 0.101 ± 0.021 |

**Note:** Values are reported as mean ± standard deviation across three random seeds. AUROC = Area Under the Receiver Operating Characteristic Curve, AUPRC = Area Under the Precision–Recall Curve, MCC = Matthews Correlation Coefficient, RF = Random Forest, LR = Logistic Regression. Bold indicates the best value in each task.

**Figure 2.**
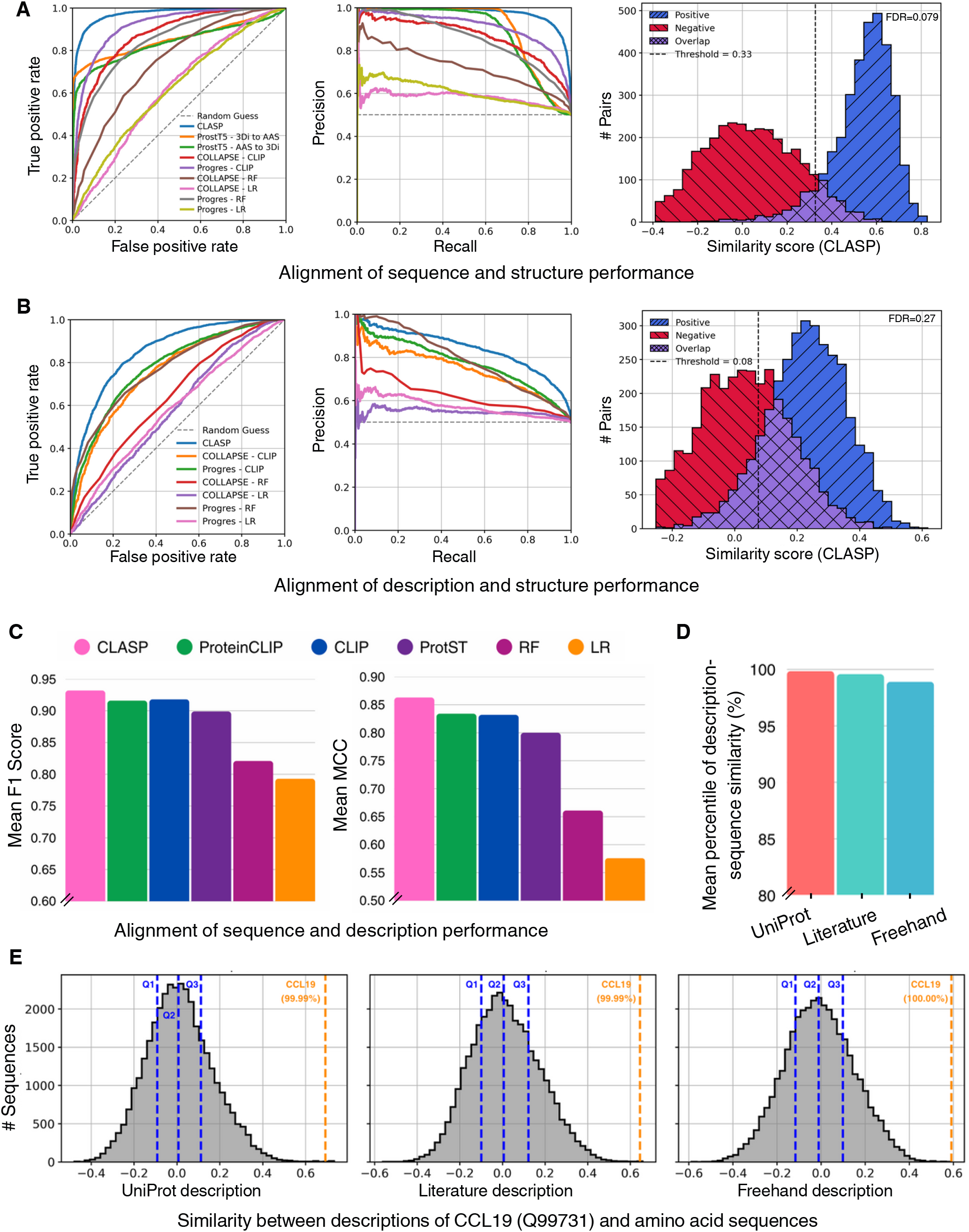
Performance of CLASP and alternative models on cross-modal classification and retrieval tasks. **(A)** ROC curves (left), PR curves (middle) and histograms of similarity-score distributions (right) for the alignment sequence and structure task for CLASP and alternative methods on a held-out test set. The dashed gray line in the ROC and the PR plots indicates the performance of random guessing. The vertical black dashed line in the histogram indicates the optimal decision threshold calculated using the validation set, with the False Discovery Rate (FDR) at that threshold indicated in the top right. Also see Table 1. **(B)** Same as (A) but for the alignment of description and structure task. **(C)** Bar plot showing the mean F1 score (left) and mean Matthews correlation coefficient (MCC, right) for the alignment of sequence and description task across three random seeds for CLASP and alternative methods on the test set. Also see Supplementary Table S1. **(D)** Bar plots showing the mean percentile rank across four proteins in sequence retrieval. Percentiles indicate the position of the correct amino acid sequence in the ranked list of 35,911 candidates. Mean percentiles are shown for UniProt (99.85% ± 0.25%), literature (99.58% ± 0.45%), and freehand (98.90% ± 1.11%) description styles. Also see Supplementary Table S2. **(E)** Distribution of CLASP similarity scores between a given description and candidate amino acid sequences for CCL19 (Q99731). Distributions are shown for UniProt (left), literature (middle), and freehand (right) description styles. Dashed blue lines indicate the quartiles of the score distribution (Q1, Q2, and Q3), and the dashed orange line indicates the position and percentile of the correct (target) protein. Also see Supplementary Table S2.

Visualization of similarity score distributions (Figure 2A, right panel) reinforces these findings. CLASP exhibits minimal overlap between positive and negative pair distributions, leading to a better separation and more precise decision boundaries. By contrast, baseline models show significant inter-class distributional overlap, indicating poorer separation of relevant and irrelevant structure–sequence pairs (Supplementary Figure S1). ROC and PR curves Figure 2A corroborate this trend, with CLASP consistently exhibiting superior performance across the score spectrum. Collectively, these results demonstrate that CLASP excels at zero-shot alignment of protein structures and sequences.

### 2.3 CLASP outperforms other models in zero-shot alignment of description and structure

Next, we set out to assess CLASP’s ability to align text-based biochemical and functional descriptions of proteins and their corresponding 3D structures using a zero-shot classification. Given a candidate description-structure pair, we computed similarity scores as the dot product between their projected embeddings in the shared space. As in the sequence-structure experiment, an optimal threshold was selected based on the validation set to maximize F1 score, and metrics were computed on a held-out test set using this threshold. Each experiment was repeated across three random splits of the dataset to ensure stability.

To serve as baseline comparisons, we substituted CLASP’s structure encoder with Progres [6] and COLLAPSE [21] embeddings and trained both CLIP-style alignment modules and traditional classifiers, Random Forest and logistic regression, on each structure–description pair.

As shown in Table 1, CLASP achieved the highest scores across all classification metrics, with a mean AUROC of 0.858 ± 0.007 and AUPRC of 0.846 ± 0.007 (also see Figure 2B). CLASP enables this accurate description-structure classification by generating embeddings whose similarity score distributions for positive and negative test pairs are well-separated; this clear separation compared to alternative models facilitates reliable thresholding and high precision during inference (Fig. 2B, Supplementary Figure S2).

### 2.4 CLASP enables sequence retrieval using different description styles

We next evaluated CLASP’s ability to align amino acid sequences and their corresponding natural language descriptions. We compared our model against ProteinCLIP [15], ProtST [16] fine-tuned using a CLIP-style contrastive objective, a standard CLIP model and two classical classifiers (RF and LR) trained on ProtT5 and BioGPT embeddings.

As shown in Figure 2C, CLASP outperformed all baselines (yet modestly), achieving the highest mean F1 score (0.932) and mean MCC (0.863). CLASP also achieved the highest mean accuracy (0.931), AUROC (0.979), and AUPRC (0.977) (Supplementary Table S1). We should point out that all tested models here (including ProteinCLIP and ProtST) utilize pLMs to obtain sequence embeddings and natural LLMs to obtain description embeddings (Methods). The rich embeddings used by these models as inputs are likely the reason why all models (even RF and LR) perform well in this task. While modest, CLASP’s consistent advantage compared to alternative models likely arises from the additional tri-modal supervision, where structural constraints implicitly regularize the relationship between sequence and description embeddings. This is particularly evident from comparing CLASP with CLIP, which uses identical description and sequence inputs and a bimodal contrastive objective instead of a tri-modal one (Figure 2C and Supplementary Table S1).

Given its good performance in aligning natural language descriptions and proteins, we next evaluated CLASP’s performance in retrieving the correct amino acid sequence from a large set of 35,911 candidate sequences, given only a textual description as a query. This evaluation has been devised to better reflect the practical utility of CLASP in real-world applications. Crucially, we designed this evaluation to probe not only retrieval performance on proteins held out from training but also robustness to variations in description types. To that end, we used three distinct types of descriptions for four proteins, each reflecting different levels of complexity: (1) curated entries from UniProt (the style which was also used during CLASP’s training phase); (2) literature-style descriptions sourced from external materials such as academic papers and Wikipedia; and (3) freehand descriptions written by a domain expert in our lab. These categories are intended to span a gradient of increasing difficulty in terms of linguistic variability and alignment with training data: UniProt entries are structured and formulaic, resembling the text seen during training; literature-style descriptions introduce more diverse phrasing and context-dependent language; and freehand inputs are the most variable, often informal and unconstrained by standard annotation guidelines. For example, the freehand description for MMP9 is: “(Matrix metalloproteinase-9) Enzyme that degrades the extracellular matrix. It is secreted by neutrophils as a component of neutrophil extracellular traps.” The full set of descriptions used in this experiment is available in Supplementary Note 1.

As can be seen in Figure 2D-E, CLASP achieved strong retrieval performance. Specifically, the corresponding sequence was ranked consistently higher than 98th percentile (Supplementary Table S2). While performance degraded with increasing linguistic complexity (as expected), CLASP retained high-rank percentiles for freehand with a mean percentile of 98.90% ± 1.11%. These results suggest that CLASP’s shared embedding space meaning-fully aligns sequences and descriptions.

### 2.5 CLASP’s embeddings cluster protein structures by family

Next, we asked whether the structure embeddings learned by the EGNN and fine tuned using CLASP’s tri-modal contrastive learning captures information regarding protein families. For this purpose, we evaluated how well CLASP’s structure embeddings cluster PDBs by their corresponding protein families. We focused on PDBs corresponding to five protein families of kinases (n = 9,997 PDBs), GPCRs (G protein-coupled receptors, n = 3,613 PDBs), ion channels (n = 1,083 PDBs), HSPs (heat shock proteins, n = 775 PDBs), and ABC transporters (n = 189 PDBs), due to their distinct functional and structural characteristics and their role in various biological processes.

Compared to Progres, ProstT5, and COLLAPSE, CLASP’s embeddings showed a better clustering performance based on the silhouette score [24], Calinski–Harabasz (CH) index [25], Kullback–Leibler (KL) divergence [26], Jensen–Shannon (JS) [27] divergence, and Davies–Bouldin index [28] (Figure 3A, Supplementary Table S3). These results suggest that embedding of PDBs corresponding to proteins of the same family are placed closer to each other in the latent space by CLASP compared to alternative models, capturing biologically meaningful patterns.

**Figure 3.**
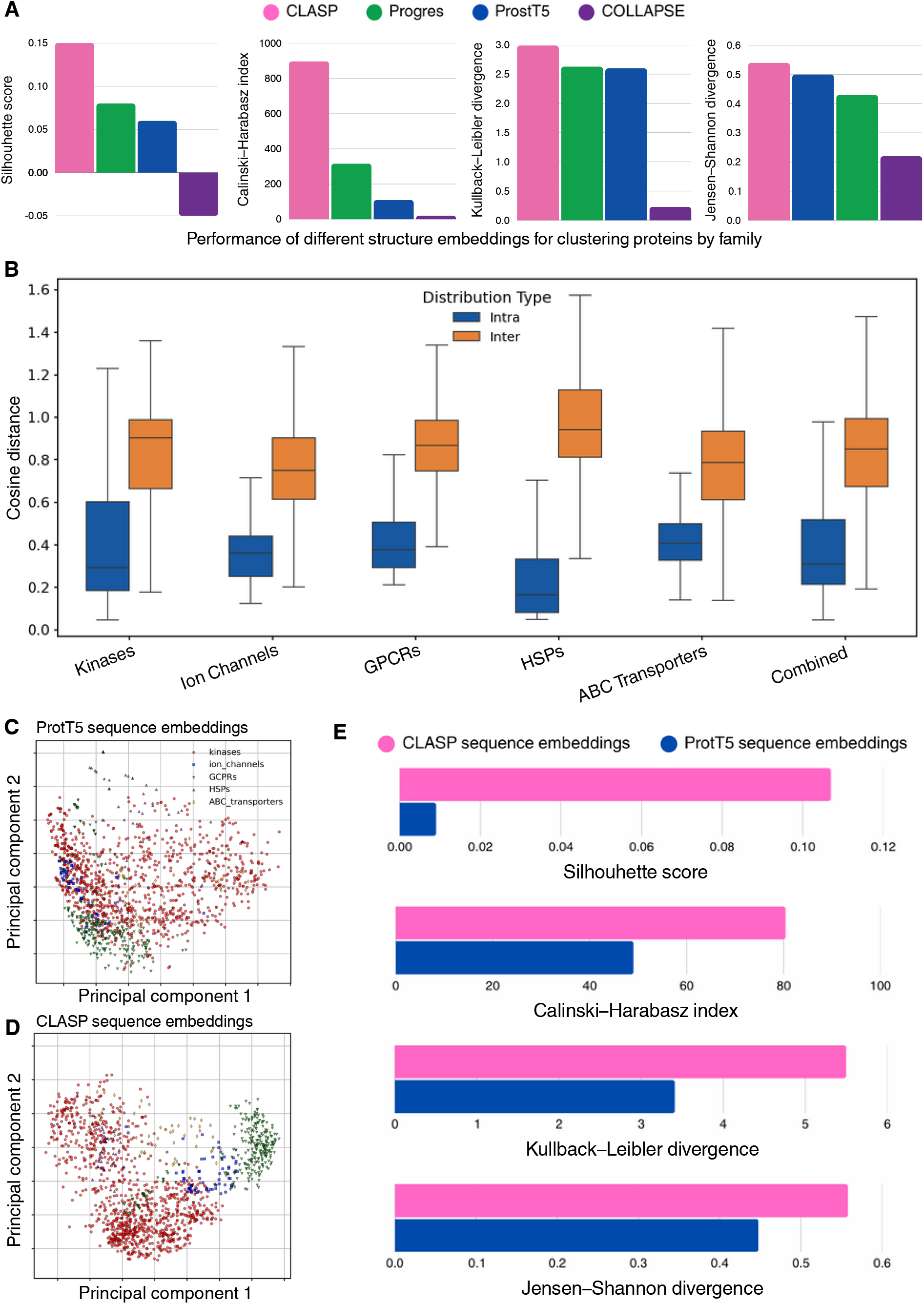
Performance of latent embeddings in clustering by protein family. **(A)** Bar plots showing the performance of structure embeddings of CLASP (pink), Progres (green), ProstT5 (blue) and COLLAPSE (purple) based on four clustering metrics: Silhouette score (far left), Calinski–Harabasz index (middle left), Kullback–Leibler divergence (middle right), and Jensen–Shannon divergence (far right). A larger value shows a better separation of clusters. **(B)** Box plots depicting the distribution of cosine distances between CLASP’s structure embeddings and a protein family centroid. Intra-cluster distances (blue) are measured from each point to its own cluster centroid, while inter-cluster distances (orange) are measured from points in other families to the same centroid. “Combined” shows the merged distribution of intra-cluster and inter-cluster distances for all families. **(C–D)** Principal component analysis (PCA) of amino acid sequence embeddings of ProtT5 (C) and CLASP (D). CLASP embeddings are those obtained after the projection layer. Each point represents an amino acid sequence and is colored and shaped by protein family: kinases (red circles), ion channels (blue squares), GPCRs (green triangles), HSPs (purple upward triangles), and ABC transporters (orange diamonds). **(E)** Bar plots comparing the clustering performance of ProtT5 sequence embeddings (blue) and CLASP sequence embeddings (pink) using the same four metrics as in panel A.

This is further confirmed by comparing the distribution of distances of embeddings of a family to their family centroid (i.e., “intra-cluster” distances) against embeddings of other families to the same centroid (“inter-cluster” distances) (Figure 3B, Supplementary Table S4). This clear separation of intra-cluster and inter-cluster distance distributions is also evident from their KL and JS divergence values, with HSPs showing the highest degree of divergence (Supplementary Table S4).

These results suggest that CLASP’s EGNN encoder combined with its tri-modal contrastive training that aligns structure embeddings with sequence and description embeddings produce functionally relevant and biologically meaningful representations with clearer family-level separation compared to other approaches.

### 2.6 Analysis of CLASP’s architecture and performance

First, we investigated whether CLASP’s amino acid sequence projection layer, which is trained to align ProtT5 sequence embeddings with structure and description modalities in a contrastive manner, enhances the separation of embeddings by protein families. Using the same five protein families as before (see Supplementary Table S5 for the number of sequence embeddings in each family), we assessed the clustering performance of the original ProtT5 embeddings and the projected CLASP embeddings.

Visually, the CLASP embeddings show a more distinct separation by protein families, based on the top two principal components, compared to the original ProtT5 embeddings (Figure 3C-D). This was also confirmed quantitatively using clustering metrics (Figure 3C-D, Supplementary Table S6). For example, the Silhouette score was approximately 12-fold higher based on CLASP embeddings, while the CH index was approximately higher by a factor of two (Figure 3E). These results suggest that the contrastive projection layer further refines the rich ProtT5 embeddings to add important, biologically-relevant information to the model’s latent representations.

Next, we performed an ablation study to assess the contribution of the EGNN and other architectural choices. First, we trained a variant of CLASP in which the EGNN was replaced with a standard graph neural network (GNN) (Table 2). On both sequence-structure and description-structure tasks, this model performed worse than CLASP, showcasing the importance of using EGNN in obtaining structure embeddings from PDBs. This improvement is primarily due to EGNN’s explicit E(3)-invariance, which ensures that learned representations depend on intrinsic geometric relationships rather than arbitrary rotations or translations of the input structure.

**Table 2.** Performance of CLASP and its variants in sequence-structure (AAS-PDB) and description-structure (DESC-PDB) alignment tasks.

| Task | Model | Accuracy | F1 Score | AUROC | AUPRC | MCC |
| --- | --- | --- | --- | --- | --- | --- |
| AAS-PDB | CLASP | **0.920 ± 0.003** | **0.920 ± 0.003** | **0.976 ± 0.002** | **0.977 ± 0.002** | **0.841 ± 0.006** |
|  | CLIP (w EGNN) | 0.905 ± 0.013 | 0.905 ± 0.014 | 0.964 ± 0.009 | 0.968 ± 0.007 | 0.810 ± 0.025 |
|  | CLASP (w GNN) | 0.844 ± 0.003 | 0.848 ± 0.001 | 0.922 ± 0.002 | 0.925 ± 0.002 | 0.688 ± 0.005 |
|  | CLASP (mean pooled) | 0.620 ± 0.169 | 0.733 ± 0.094 | 0.642 ± 0.209 | 0.645 ± 0.207 | 0.238 ± 0.341 |
|  | CLASP (max pooled) | 0.610 ± 0.023 | 0.709 ± 0.009 | 0.679 ± 0.039 | 0.642 ± 0.047 | 0.300 ± 0.035 |
| DESC-PDB | CLASP | **0.770 ± 0.011** | **0.791 ± 0.005** | **0.858 ± 0.007** | **0.846 ± 0.007** | **0.552 ± 0.018** |
|  | CLIP (w EGNN) | 0.764 ± 0.022 | 0.783 ± 0.024 | 0.840 ± 0.024 | 0.814 ± 0.019 | 0.538 ± 0.047 |
|  | CLASP (w GNN) | 0.686 ± 0.008 | 0.728 ± 0.006 | 0.774 ± 0.009 | 0.756 ± 0.010 | 0.391 ± 0.016 |
|  | CLASP (max pooled) | 0.598 ± 0.011 | 0.700 ± 0.002 | 0.661 ± 0.012 | 0.621 ± 0.015 | 0.270 ± 0.011 |
|  | CLASP (mean pooled) | 0.569 ± 0.098 | 0.691 ± 0.034 | 0.597 ± 0.136 | 0.593 ± 0.124 | 0.143 ± 0.202 |

**Note:** Values are reported as mean ± standard deviation across three random seeds. AUROC = Area Under the Receiver Operating Characteristic Curve, AUPRC = Area Under the Precision-Recall Curve, MCC = Matthews Correlation Coefficient. Bold indicates the best value in each column.

We further investigated the effect of the graph-level aggregation strategy by replacing sum pooling with mean and max pooling, to assess whether CLASP’s performance could be biased by protein chain length. As shown in Table 2, both mean and max pooling led to substantial and consistent performance degradation across sequence– structure and description–structure alignment tasks, with notably lower MCC and AUPRC values. These results support sum pooling as the most effective aggregation strategy for CLASP’s protein graphs.

Next, we examined the effect of weighting the tri-modal contrastive loss terms, which are equally weighted in the default CLASP configuration. We retrained CLASP under nine additional loss-weighting schemes spanning asymmetric and single-modality pairings and evaluated performance across all three alignment tasks (Supplementary Table S7). We observed that, perhaps unsurprisingly, optimal weightings differ by task, reflecting differences in signal strength across modality pairs. However, when aggregating performance by ranking configurations using MCC across tasks, the balanced configuration used in the main paper, while not always the top performer for any single task, achieves the best overall mean rank (Supplementary Table S8), supporting equal loss weighting as a robust compromise across modalities.

We also assessed CLASP’s sensitivity to the contrastive temperature parameter τ. We evaluated the default value of τ = 0.07 (which was adopted from the original CLIP paper [14]) via a sweep over a broad range of temperatures (Supplementary Figure S3). Performance was stable at lower temperatures (τ < 0.1) across all alignment tasks, while higher values led to degraded performance, albeit minor, indicating that CLASP is not overly sensitive within a reasonably low-temperature regime and supporting our choice of τ = 0.07.

Next, we examined whether CLASP’s gains could be driven by trivial lexical cues in the description modality, such as explicit protein names. To this end, we conducted an ablation in which protein names were removed from all natural language descriptions during training and evaluation. As shown in Supplementary Tables S9–S11, removing names results in only modest and consistent performance decreases across description-structure alignment, sequence–description alignment, and large-scale sequence retrieval tasks. In particular, retrieval mean percentile ranks remain above 98% across all description styles (Supplementary Table S11), indicating that CLASP’s cross-modal alignment is not dependent on exact name matching but instead reflects learnt semantically meaningful relationships.

To assess the role of tri-modal contrastive training, we trained two variations of CLASP, in which the original EGNN encoder was used, but the tri-modal objective function was replaced by a two-modality CLIP objective. Both models showed an inferior performance compared to CLASP in their respective tasks of structure-sequence or structure-description alignment (Table 2). Interestingly, a larger drop in the performance was observed when we replaced the EGNN with GNN compared to when we changed the tri-modal objective function, further emphasizing the importance of structure encoder in CLASP’s performance.

Finally, we asked whether insights could be obtained by characterizing cases in which CLASP succeeded, but CLIP did not. To quantify this, we computed the Skill-Normalized Concordance (SNC) [3] between the predictions of these models across the zero-shot classification tasks, observing only moderate agreement between these methods (sequence-structure: 0.53 ± 0.05; description-structure: 0.57 ± 0.03). An analysis of proteins correctly classified by CLASP but not by CLIP using two-sided Mann–Whitney U tests revealed no significant difference between their sequence lengths compared to the remainder of the test set (sequence-structure: p=0.56, description-structure: p=0.30). Similarly, CLASP-only successes did not show enrichment in any organism, UniProt keywords, or functional annotations (hypergeometric test followed by Benjamini–Hochberg correction; Supplementary Tables S12–S13). These results suggest that CLASP’s gains over CLIP are not driven by trivial sequence-length differences or by overrepresentation of any specific protein subset or functional class, but instead reflect a broader, distributed advantage consistent with the additional structural inductive bias introduced by incorporating explicit 3D information.

## 3 Discussion

CLASP is a unified tri-modal framework that integrates protein structure, sequence, and natural language description into a shared representation space through contrastive pretraining. By jointly aligning structural embeddings generated by an E(3)-invariant graph neural network with precomputed amino acid sequence and language embeddings, CLASP captures complementary views of a protein and significantly improves cross-modal alignment tasks without requiring task-specific supervision.

The tri-modal contrastive learning framework of CLASP operates under the assumption that embeddings from distinct modalities can be aligned in a shared latent space. Here, these modalities correspond to sequence (discrete 1D strings), structure (continuous 3D geometric objects), and functional descriptions (linguistic abstractions) that are ontologically distinct with non-isomorphic intrinsic similarity metrics. Despite these differences, this alignment is theoretically grounded in the causal structure of protein biology: sequence determines structure, and structure largely determines function, implying substantial mutual information across modalities. The shared embedding space can thus be understood as capturing the invariant aspects of protein identity that are expressed differently across modalities. Empirically, our clustering analysis and cross-modal retrieval results demonstrate that the learned shared space is not arbitrary but preserves biologically meaningful relationships, supporting the validity of this modeling approach. Nevertheless, it is important to note that cross-modal alignment also serves as a practical co-regularization strategy that may enhance representation quality even in the absence of perfect geometric commensurability.

We demonstrated that CLASP enables accurate zero-shot classification across all modality pairs, outperforming baselines on symmetric sequence-structure, description-structure, and sequence-description alignment tasks. In particular, its significant improvement compared to its own variations in which only two data modalities were used along with a CLIP objective function signifies the added value of a joint tri-modal contrastive training to obtain rich embeddings.

We showed that CLASP also enables sequence retrieval from over 35,000 candidates using descriptions written in multiple styles (including freehand summaries, curated UniProt entries, and literature-style prose) with mean retrieval percentiles above 98% across all categories. This robustness to language variability reflects the model’s ability to generalize beyond the structured training data, suggesting potential applications in semantic protein search engines or literature-driven protein design.

In addition to its cross-modal alignment capabilities, CLASP produces embeddings that better reflect protein family structure. Structural embeddings obtained from CLASP cluster into functionally coherent groups, as demonstrated by various clustering metrics. We hypothesize that this effect arises in part from the hierarchical nature of the EGNN encoder: early message-passing layers capture local geometric and physicochemical interactions encoded by node features and distance-based edges, while successive layers integrate longer-range topological and geometric relationships across the graph, yielding embeddings that reflect both local residue environments and global structural organization. These trends hold for both structure and sequence embeddings, suggesting that CLASP’s contrastive objective encourages alignment not only across modalities but also within each individual modality. The inclusion of natural language descriptions during training appears to act as a functional regularizer, encouraging structural and sequential encoders to capture biologically meaningful features that reflect known family-level distinctions. This multi-modal supervision leads to more discriminative and functionally informative representations, outperforming structure-only baselines on all clustering metrics.

An additional consideration is whether the modality-specific linear projection layers preserve biologically meaningful structure in the original embeddings. While there is no formal guarantee that linear projections fully preserve modality-specific topology, the projections used in CLASP are intentionally low-capacity and linear, limiting them to bounded transformations of already informative representations. Empirically, the preservation of biological meaning is supported by improved protein family clustering after projection relative to raw ProtT5 embeddings, as well as strong zero-shot alignment and retrieval performance across modalities. Together, these results suggest that the projection layers align modalities without destroying underlying biophysical or functional structure.

Ablation studies confirmed that both the tri-modal training objective and the E(3)-invariant architecture are essential to CLASP’s performance. Replacing the EGNN with a standard GNN reduces MCC by over 15 points, while training with a bi-modal CLIP-style objective leads to consistently worse performance in both sequence-structure and description-structure tasks. These findings emphasize that CLASP’s architecture gains do not come from any single component in isolation but from the interaction between its geometry-aware encoder and its tri-modal alignment objective. The symmetry and shared supervision across all modality pairs are particularly important, as they allow information from one modality (such as descriptive language) to inform representations in another (such as structure), resulting in a unified embedding space with enhanced generalization capacity.

CLASP does, however, face limitations. The coverage of proteins with structural data in the training dataset is constrained by the availability of high-confidence PDB entries, which can introduce bias against certain protein classes, such as intrinsically disordered regions or membrane proteins. The model also operates with frozen pLM and LLM backbones (ProtT5 and BioGPT, respectively), and although this reduces computational overhead, it may limit deeper cross-modal synergy; however, representations from all three modalities are still passed through trainable linear projection layers, enabling alignment to be learned in the shared latent space despite frozen backbones (as noted in the sequence–description task). Joint fine-tuning or adapter-based approaches could further enhance performance, but require careful calibration to prevent overfitting or catastrophic forgetting [29].

Despite these constraints, CLASP establishes a strong foundation for tri-modal protein representation learning and opens new directions for multi-modal biological modeling. By embedding structure, sequence, and natural language into a single, biologically grounded space, CLASP unifies molecular and semantic information, enabling tasks that bridge mechanistic and functional domains. The model’s ability to align and retrieve proteins across modalities (without retraining) suggests applications in protein annotation, drug discovery, and automated literature synthesis. Future extensions may incorporate additional modalities such as evolutionary context or tissue-specific expression, and conditional generation models may allow users to synthesize candidate structures from textual prompts or functional targets.

## 4 Methods

### 4.1 Data acquisition and preprocessing

The main CLASP dataset consists of *n* = 35, 911 unique UniProt Knowledgebase (UniProtKB) [8] accession numbers (UPKB ACs), each linked to at least one Protein Data Bank (PDB) entry. For each UPKB AC, we collected the corresponding structural information, including PDB ID, chain identifier, raw chain sequence, and annotated residue start and end positions. On average, each protein was associated with 8.61 PDB entries (median = 2), reflecting structural variation due to conformational changes or crystallization conditions. These associations were extracted programmatically from the reviewed (Swiss-Prot) UniProtKB XML archive (https://www.uniprot.org/downloads) using custom parsers. Although the resulting dataset is modest in size compared to those used to train large protein language models, CLASP is intentionally designed with a relatively small number of trainable parameters; in this regime, training on a curated, high-confidence PDB-derived dataset is appropriate and advantageous, as it leverages experimentally validated structures while mitigating overfitting risk.

To control for sequence homology across dataset splits, proteins were first clustered based on amino acid sequence similarity before train–validation–test partitioning. Specifically, proteins were clustered using MMSeqs2 [30], with any two proteins sharing ≥30% sequence identity assigned to the same cluster. Training (80%), validation (10%), and testing (10%) splits were then constructed at the cluster level, ensuring that all proteins belonging to the same sequence-similarity cluster were assigned to the same split. This split was repeated independently across three random seeds to evaluate stability and reproducibility. For each training set associated with a seed, we generated five variations, each containing a different randomly selected PDB entry for the training proteins (when multiple PDB entries were available). This was done to introduce structural diversity during training. During training, we cycled through these five sets across epochs, ensuring that each protein was paired with a distinct structure every fifth epoch, thereby enriching the training distribution.

For evaluation, we constructed the same number of positive and negative pairs from each test set, resulting in a total of 7276, 7212, and 6868 pairs for the three seeds. For tasks involving structure, each positive pair was created by selecting a protein sequence or description and randomly sampling one of its associated PDBs. A corresponding negative pair was formed by pairing the same sequence or description with a PDB belonging to a different protein from the test set. For sequence-description tasks, the same approach was used except that the negative pair was generated by sampling a description associated with a different protein in the test set. The same data splits were used across all models to ensure a fair comparison.

For the sequence modality, we used 1024-dimensional per-protein amino acid embeddings generated by the ProtT5-XL-UniRef50 model [2], a protein language model trained using a masked language modeling objective on large-scale protein sequence databases. These embeddings were directly downloaded from the UniProt website’s embedding repository at https://www.uniprot.org/help/embeddings. The provided embeddings are pre-computed by the UniProt consortium using the bio_embeddings framework, specifically the prottrans_t5_xl_u50 model. Only canonical sequences from the UniProtKB/Swiss-Prot database were retained. Importantly, sequence and structure inputs are sourced independently, as sequences are not derived from PDB files.

For the natural language modality, protein descriptions were curated from UniProt’s structured annotations, specifically the “Protein names” and “Function” fields [8]. These text descriptions included formal protein names, synonyms, and functional summaries. Descriptions were tokenized and embedded using BioGPT [20], a transformer-based language model fine-tuned on biomedical corpora. We used the official HuggingFace implementation (https://huggingface.co/microsoft/BioGPT) to generate 1024-dimensional embeddings for each description. For the sequence retrieval task, the protein descriptions (whether UniProt, Literature, or Freehand styles) were processed in a similar manner.

### 4.2 Protein structure graph representation

In CLASP, protein structures are represented as graphs, enabling the model to capture intricate geometric and biochemical information. These graphs are constructed using Graphein [19], a specialized library for protein structural data. Each graph encodes atoms as nodes, annotated with biochemical and spatial features, and edges representing spatial relationships or biochemical interactions. Graphs were generated using Graphein’s default “ProteinGraphConfig”, with no custom parameters modified during construction.

Proteins were matched with their corresponding three-dimensional structures using the Uniprot SWISS-PROT Knowledgebase. More precisely, the Uniprot knowledgebase contains information linking proteins to Protein Data Bank (PDB) files in which they are present, as well as the relevant polypeptide chain. Graphein was used to download PDB files, construct structure graphs from them, and filter all but the relevant polypeptide chains.

Each node in the graph was annotated with seven biochemical features corresponding to the Meiler descriptors as implemented in Graphein: hydrophobicity, steric bulk (volume), polarity, charge, aromaticity, flexibility, and hydrogen-bonding propensity [31]. These descriptors offer a compact yet comprehensive characterization of the biochemical nature of each residue. Additionally, the 3-dimensional (3D) spatial coordinates of each node are preserved, providing geometric information.

Edges in the graph were defined based on spatial proximity, capturing relationships between nodes within a certain distance threshold. Covalent and non-covalent interactions were not treated as separate edge types but are implicitly encoded through Euclidean inter-atomic distances. Each edge was weighted by the Euclidean distance between the 3D coordinates of its connected nodes, reflecting the protein’s spatial arrangement. These weights are essential for tasks involving geometric analysis, as they allow the model to consider both local and global structural relationships.

These graphs were converted into PyTorch Geometric Data objects for integration with the downstream encoder. The nodes were indexed to ensure consistency, and tensors were generated for node features, edge indices, edge attributes (distances), and node positions (coordinates). This ensures that the graphs are optimized for efficient processing in the EGNN.

### 4.3 E(3)-invariant graph neural network architecture

Graph neural networks (GNNs) are a class of deep learning models that operate on graph-structured data, allowing information to propagate along edges between nodes. In molecular settings, GNNs are commonly used to model interactions between atoms or molecules, where the graph topology reflects the spatial or chemical connectivity of the nodes. However, conventional GNNs typically ignore the geometric symmetries of 3D space, making them suboptimal for tasks where spatial configuration is biologically meaningful.

To address this limitation, CLASP employs an E(3)-invariant graph neural network (EGNN) [12] that incorporates spatial geometry while remaining insensitive to the protein’s absolute orientation or position in space. This invariance is achieved by encoding pairwise squared Euclidean distances between nodes, quantities that are mathematically invariant under rotations, translations, and reflections in 3D. These distance-based edge features enable the model to capture biologically relevant internal geometric relationships while discarding extraneous coordinate information. Such invariance ensures that structurally identical proteins yield identical embeddings [12]. Our implementation closely follows the implementation described in TeachOpenCADD [32].

The EGNN begins by transforming node features into an initial embedding space. Each node *i* in the graph has features **f**_*i*_ ∈ ℝ^*k*^ (here *k* = 7), which are mapped to an embedding 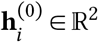 using a learnable linear transformation

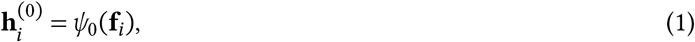

where *ψ*_0_ is a linear projection. A central component of the EGNN is its invariant message-passing mechanism, which iteratively updates node embeddings based on their relationships with neighboring nodes. At each layer *l*, pairwise distances 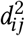 between nodes *i* and *j* are computed from their 3D coordinates **c**_*i*_ and **c**_*j*_

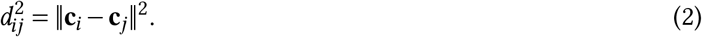

These distances serve as edge attributes, capturing geometric relationships between nodes. Using the embeddings of the source node 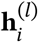 and target node 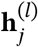, along with the distance 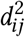, the model computes a message **m**_*ij*_ through a learnable function *ϕ*_*l*_. The messages from all neighbors of a node are aggregated via summation, yielding the total message for node *i*:

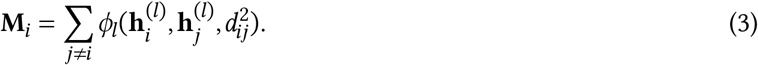

This aggregated message **M**_*i*_ is combined with the node’s current embedding 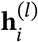 to compute the updated embedding 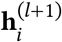. A residual connection ensures stable training and preserves information from earlier layers according to.

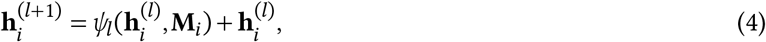

where *ψ*_*l*_ is a learnable multi-layer perceptron (MLP) that integrates the node embedding with the aggregated message.

After *L* message-passing layers (here we used *L* = 3), which is sufficient for dense, radius-based protein graphs where each layer aggregates information from a large spatial neighborhood [12, 33], the final node embeddings 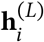 are aggregated using a sum pooling operation to produce a graph-level embedding. This embedding is then processed through a two-layer feedforward neural network *ψ*_readout_, defined as

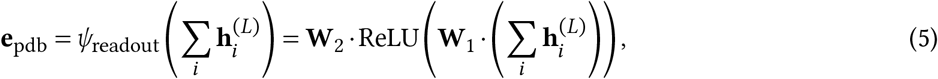

where **W**_1_ and **W**_2_ are learnable weight matrices corresponding to linear layers. All non-linearities throughout the model, including the initial embedding layer *ψ*_0_, the message and update functions *ϕ*_*l*_ and *ψ*_*l*_, and the readout function *ψ*_readout_, use the ReLU activation function. Residual connections are employed in each message-passing layer to stabilize training and facilitate the flow of gradients.

### 4.4 Alignment module architecture - 3D CLIP

The Contrastive Language-Image Pretraining (CLIP) model [14] provides a mechanism for aligning representations from two distinct modalities (typically images and text) into a shared embedding space. A central aspect of CLIP’s training procedure is its employment of contrastive learning, where the embeddings of matched pairs are “pulled” closer together, and those of mismatched pairs are “pushed” apart. This is to say that the distance between pairs of embeddings which match the same protein are minimized, while the distance of mismatched pairs are maximized. This framework enables tasks such as cross-modal retrieval and zero-shot classification by learning to map semantically related content closer in the embedding space [14]. The CG3D model (CLIP goes 3D) [18] extends this approach by introducing a dedicated 3D encoder trained to align point cloud representations with CLIP’s image and text embeddings, thereby enabling zero-shot understanding directly from geometric data. This extension yields significant performance benefits in 3D recognition, retrieval, and scene querying tasks [18]. In CLASP, the CG3D methodology has been adapted to align protein natural language descriptions, amino acid sequences, and structures (PDBs). The model first processes each modality independently through a modality-specific linear projection layer. These projection heads map each input vector to a unified 512-dimensional embedding space. Formally, let **e**_pdb_, **e**_aas_, **e**_desc_ be the input embeddings for a given protein’s structure (from the EGNN), sequence (from ProtT5), and description (from BioGPT), respectively. These are first transformed via independently parameterized modality-specific linear projection layers and then *L*^2^-normalized to unit length, represented as **x**_pdb_, **y**_aas_, and **z**_desc_ (**x, y, z**, for brevity) (Figure 1B-C).

Given a batch of *n* proteins, three pairwise cross-modality similarity matrices **S**_mod1, mod2_ ∈ ℝ^*n*×*n*^ are formed. Specifically, the (*i, j*)-th entry of each such matrix captures the cross-modality similarity of protein *i* and protein *j*, calculated as

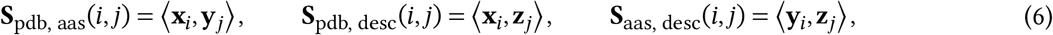

where ⟨⋅, ⋅⟩ denotes the dot product (Figure 1C). Importantly, 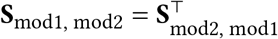, meaning that similarity is evaluated symmetrically across modalities rather than directionally. These similarity matrices are used to calculate the loss function during training and serve as the foundation for downstream tasks during inference such as cross-modal retrieval and classification.

### 4.5 Training process and loss function

CLASP is trained end-to-end using a tri-modal contrastive learning objective inspired by the CG3D framework [18], extended to protein data. The core idea is to align embeddings from three modalities - structure, sequence, and description - by maximizing similarity between matched pairs and minimizing similarity between mismatched pairs. For each modality pair, a symmetric cross-entropy loss is computed across the pairwise similarity matrices **S**_pdb, aas_, **S**_pdb, desc_, and **S**_aas, desc_. Specifically,

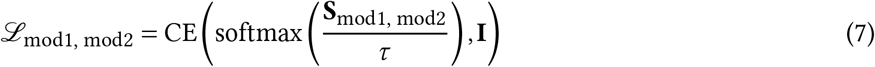

where mod1, mod2 ∈ {pdb, aas, desc}, τ is a temperature parameter controlling the sharpness of the softmax distribution, CE is the cross-entropy, and **I** is the identity matrix representing the correct alignment between corresponding pairs in the batch. The cross-entropy loss encourages the diagonal entries of the similarity matrix (representing correctly matched pairs) to have the highest similarity scores, while suppressing scores of mismatched (off-diagonal) pairs.

The total contrastive loss is computed as the average of the pairwise losses across all modality pairs. To prevent overfitting and encourage generalization, an *L*^2^ regularization term is applied to the parameters of the structure encoder. The final objective is

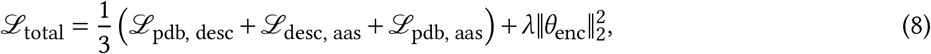

where *θ*_enc_ denotes the encoder parameters, and *λ* is the regularization weight. Training is performed with the Adam optimizer using early stopping based on validation loss. Hyperparameters (*τ* and *λ*) were selected based on performance on the held-out validation set. Models are trained across three distinct random seeds to ensure reproducibility and robustness.

### 4.6 Baseline methods

We chose two state of the art PDB embedding methods to serve as baseline comparisons: Progres [6], which generates structure-aware protein embeddings by encoding local atomic environments using a hierarchical graph convolutional network and contrastive learning on protein graph pairs; and COLLAPSE [21], a self-supervised framework that learns local structure-function representations by aligning residue-centered 3D environments across homologous proteins using evolutionary conservation signals from multiple sequence alignments. To assess structure alignment quality, we trained separate alignment modules on these fixed structure embeddings using the same amino acid sequence and natural language description embeddings employed in CLASP. For each structure embedding method (Progres [6] and COLLAPSE [21]), we trained a standard CLIP [14] (including trainable projection layers) and two conventional classifiers, Random Forest and logistic regression, trained on concatenated structure–sequence or structure–description embedding pairs. All models were evaluated on the same training, validation, and test splits, ensuring a fair comparison. Progres and COLLAPSE were selected because they explicitly encode 3D geometry, making them well-suited baselines for structure-related alignment tasks.

In addition to Progres and COLLAPSE, we included ProstT5 [22] as a baseline. ProstT5 is a bilingual protein language model that extends ProtT5 by learning a joint representation of amino acid sequences and discrete structural 3Di sequences, enabling translation between sequence and structure. We evaluated ProstT5 on the sequence-structure alignment task by leveraging its generative capabilities in both directions (3Di→AAS and AAS→3Di). Specifically, PDB structures were first converted into 3Di sequences using Foldseek [34], after which pretrained ProstT5 was used to generate the corresponding target modality. For each candidate pair, we computed a Smith–Waterman alignment score [23] between the generated output and the ground-truth target sequence. As with CLASP and all CLIP-based baselines, an optimal decision threshold was selected on the validation set to maximize the F1 score and then applied to the held-out test set. This procedure was repeated independently for both translation directions and across all random seeds, yielding two ProstT5 baselines for sequence-structure alignment.

For sequence-description alignment tasks, we considered two alternative baselines: ProteinCLIP [15], which contrasts amino acid sequences with functional text using frozen pretrained language models and lightweight projection heads; and ProtST [16], a multi-modal framework that jointly trains protein and biomedical text representations using a combination of masked language modeling and cross-modal alignment tasks. We retrained ProteinCLIP [15] on our data split using their ProtT5 [2] sequence and OpenAI text-embedding-3-large (https://platform.openai.com/docs/models/text-embedding-3-large) description embeddings. For ProtST [16], we fine-tuned a CLIP-style [14] alignment model on their zero-shot classification setup by projecting their ESM-1b [35] sequence embeddings and PubMedBERT [36] description embeddings into a shared embedding space using a contrastive InfoNCE loss. Finally, we trained a standard CLIP [14] model (with trainable projection layers) and two conventional classifiers - Random Forest and logistic regression (on concatenated sequence-description embedding pairs) - using the same frozen BioGPT [20] and ProtT5 [2] embeddings as in CLASP. All models were evaluated on the same training, validation, and test splits to ensure a fair comparison.

### 4.7 Implementation details

CLASP was implemented in the Python programming language using the PyTorch deep learning framework. Training was performed for up to 500 epochs using the Adam optimizer (learning rate 0.001), batch size 8, and early stopping with a patience of 40. All models were trained and evaluated on an NVIDIA RTX 3090 GPU with 32 CPU cores. Each seed-specific run took approximately 13.08 hours (∼2 minutes per epoch). The structure encoder used in CLASP was an EGNN with 3 message passing layers, 7 input node features, hidden dimensionality of 16, and output embedding size of 512. Final graph-level representations were produced using sum pooling followed by a two-layer MLP with ReLU activations. Tri-modal supervision was applied using a multi-modal contrastive loss with a temperature of 0.07 and L2 regularization (λ = 0.01) on the EGNN parameters.

### 4.8 Clustering evaluation and metrics

Quantitative metrics include the silhouette score (measuring cohesion versus separation using cosine distances where higher is better) [24], Davies–Bouldin index (assessing average cluster similarity where lower is better) [28], and Calinski–Harabasz index (ratio of between- to within-cluster dispersion where higher is better) [25]. Additionally, we calculated intra-cluster cosine distances from each point to its family centroid and inter-cluster distances from points of other families to the same centroid; we evaluated distributional divergence between intra- and inter-cluster distances with Kullback–Leibler [26] and Jensen–Shannon [27] divergences computed on normalized histograms.

## Data Availability

All datasets and embeddings used in the analyses herein are made available under a the Creative Commons Attribution-NonCommercial-ShareAlike 4.0 International license. Data can be accessed via Internet Archive: https://archive.org/details/clasp_data

## Code Availability

The CLASP source code is made available under the GNU Affero General Public License v3. The code-base can be accessed via GitHub: https://github.com/Emad-COMBINE-lab/clasp.

## Supplementary Information

Except for Supplementary Tables S12 and S13 that are provided as separate files, all supplementary information, including Supplementary Tables, Supplementary Figures, and Supplementary Notes are provided in Supplementary File S1.

## Acknowledgments

This work was supported by grants from Natural Sciences and Engineering Research Council of Canada (NSERC) [RGPIN-2019-04460] (A.E.) and Canada Foundation for Innovation (CFI) JELF [project 40781]. This research was enabled in part by support provided by Calcul Québec (www.calculquebec.ca) and the Digital Research Alliance of Canada (alliancecan.ca). We acknowledge the support of Orsolya Lapohos for generating the freehand protein descriptions employed in the sequence retrieval evaluation.

