## Supplementary File S1 for "Multi-Modal Protein Representation Learning with CLASP"

### 1 Supplementary Notes

#### 1.1 Full set of natural language descriptions used in retrieval tasks

TAP2 (Q03519)

**UniProt:** "PROTEIN_NAMES: Antigen peptide transporter 2, ATP-binding cassette sub-family B member 3, Peptide supply factor 2, Peptide transporter PSF2, Peptide transporter TAP2, Peptide transporter involved in antigen processing 2, Really interesting new gene 11 protein. PROTEIN_DESCRIPTION: ABC transporter associated with antigen processing. In complex with TAP1 mediates unidirectional translocation of peptide antigens from cytosol to endoplasmic reticulum (ER) for loading onto MHC class I (MHCI) molecules (, ). Uses the chemical energy of ATP to export peptides against the concentration gradient (). During the transport cycle alternates between 'inward-facing' state with peptide binding site facing the cytosol to 'outward-facing' state with peptide binding site facing the ER lumen. Peptide antigen binding to ATP-loaded TAP1-TAP2 induces a switch to hydrolysis-competent 'outward-facing' conformation ready for peptide loading onto nascent MHCI molecules. Subsequently ATP hydrolysis resets the transporter to the 'inward facing' state for a new cycle (, , ). Typically transports intracellular peptide antigens of 8 to 13 amino acids that arise from cytosolic proteolysis via IFNG-induced immunoproteasome. Binds peptides with free N- and C-termini, the first three and the C-terminal residues being critical. Preferentially selects peptides having a highly hydrophobic residue at position 3 and hydrophobic or charged residues at the C-terminal anchor. Proline at position 2 has the most destabilizing effect (, , ). As a component of the peptide loading complex (PLC), acts as a molecular scaffold essential for peptide-MHCI assembly and antigen presentation (, , )" ([1]; `www.uniprot.org/uniprotkb/Q03519/entry`)

**Literature:** "Tap2 (also known as antigen peptide transporter 2), which is an eukaryotic protein belonging to the ABC transporter family. It plays a crucial role in the processing and presentation of the MHC class I-restricted antigens. It is a half transporter that forms a complex with Tap1. This complex translocates antigens from the cytoplasm to the endoplasmic reticulum for loading onto MHC class I molecules" ([2]; `www.ebi.ac.uk/interpro/entry/InterPro/IPR005293/`)

**Freehand:** "(Antigen peptide transporter 2) A transporter associated with the endoplasmic reticulum. Assists with intracellular peptide loading onto major histocompatibility complex class I for antigen presentation to T cells."

COX2 (P00405)

**UniProt:** "PROTEIN_NAMES: Cytochrome c oxidase subunit 2, Cytochrome c oxidase polypeptide II. PROTEIN_DESCRIPTION: Component of the cytochrome c oxidase, the last enzyme in the mitochondrial electron transport chain which drives oxidative phosphorylation. The respiratory chain contains 3 multisubunit complexes succinate dehydrogenase (complex II, CII), ubiquinol-cytochrome c oxidoreductase (cytochrome b-c1 complex, complex III, CIII) and cytochrome c oxidase (complex IV, CIV), that cooperate to transfer electrons derived from NADH and succinate to molecular oxygen, creating an electrochemical gradient over the inner membrane that drives transmembrane transport and the ATP synthase. Cytochrome c oxidase is the component of the respiratory chain that catalyzes the reduction of oxygen to water. Electrons originating from reduced cytochrome c in the intermembrane space (IMS) are transferred via the dinuclear copper A center (CU(A)) of subunit 2 and heme A of subunit 1 to the active site in subunit 1, a binuclear center (BNC) formed by heme A3 and copper B (CU(B)). The BNC reduces molecular oxygen to 2 water molecules using 4 electrons from cytochrome c in the IMS and 4 protons from the mitochondrial matrix" ([1]; `www.uniprot.org/uniprotkb/P00405/entry`)

**Literature:** "Cytochrome c oxidase (7.1.1.9) [1, 2] is an oligomeric enzymatic complex which is a component of the respiratory chain and is involved in the transfer of electrons from cytochrome c to oxygen. In eukaryotes this enzyme complex is located in the mitochondrial inner membrane; in aerobic prokaryotes it is found in the plasma membrane. The enzyme complex consists of 3-4 subunits (prokaryotes) to up to 13 polypeptides (mammals)." ([2]; `www.ebi.ac.uk/interpro/entry/InterPro/IPR011759/`)

Subunit 2 (CO II) transfers the electrons from cytochrome c to the catalytic subunit 1. It contains two adjacent transmembrane regions in its N terminus and the major part of the protein is exposed to the periplasmic or to the mitochondrial intermembrane space, respectively. CO II provides the substrate-binding site and contains a copper centre called Cu(A) (see IPR001505), probably the primary acceptor in cytochrome c oxidase. An exception is the corresponding subunit of the cbb3-type oxidase which lacks the copper A redox-centre. Several bacterial CO II have a C-terminal extension that contains a covalently bound haem c."

**Freehand:** "(Cytochrome c oxidase subunit 2) Mitochondrial enzyme that participates in the electron transport chain."

<u>MMP9 (P14780)</u>

**UniProt:** "PROTEIN_NAMES: Matrix metalloproteinase-9, 92 kDa gelatinase, 92 kDa type IV collagenase, Gelatinase B. PROTEIN_DESCRIPTION: Matrix metalloproteinase that plays an essential role in local proteolysis of the extracellular matrix and in leukocyte migration (, , ). Could play a role in bone osteoclastic resorption (By similarity). Cleaves KiSS1 at a Gly-|-Leu bond (). Cleaves NINJ1 to generate the Secreted ninjurin-1 form (). Cleaves type IV and type V collagen into large C-terminal three quarter fragments and shorter N-terminal one quarter fragments (). Degrades fibronectin but not laminin or Pz-peptide" ([1]; `www.uniprot.org/uniprotkb/P14780/entry`)

**Literature:** "Matrix metalloproteinase (MMP)-9, one of the most widely investigated MMPs, regulates pathological remodeling processes that involve inflammation and fibrosis in cardiovascular disease. MMP-9 directly degrades extracellular matrix (ECM) proteins and activates cytokines and chemokines to regulate tissue remodeling. MMP-9 deletion or inhibition has proven overall beneficial in multiple animal models of cardiovascular disease." [3]

**Freehand:** "(Matrix metalloproteinase-9) Enzyme that degrades the extracellular matrix. It is secreted by neutrophils as a component of neutrophil extracellular traps"

<u>CCL19 (Q99731)</u>

**UniProt:** "PROTEIN_NAMES: C-C motif chemokine 19, Beta-chemokine exodus-3, CK beta-11, Epstein-Barr virus-induced molecule 1 ligand chemokine, Macrophage inflammatory protein 3 beta, Small-inducible cytokine A19. PROTEIN_DESCRIPTION: May play a role not only in inflammatory and immunological responses but also in normal lymphocyte recirculation and homing. May play an important role in trafficking of T-cells in thymus, and T-cell and B-cell migration to secondary lymphoid organs. Binds to chemokine receptor CCR7. Recombinant CCL19 shows potent chemotactic activity for T-cells and B-cells but not for granulocytes and monocytes. Binds to atypical chemokine receptor ACKR4 and mediates the recruitment of beta-arrestin (ARRB1/2) to ACKR4" ([1]; `www.uniprot.org/uniprotkb/Q99731/entry`)

**Literature:** "Chemokine (C-C motif) ligand 19 (CCL19) is a protein that in humans is encoded by the CCL19 gene.[5][6]

This gene is one of several CC cytokine genes clustered on the p-arm of chromosome 9. Cytokines are a family of secreted proteins involved in immunoregulatory and inflammatory processes. The CC cytokines are proteins characterized by two adjacent cysteines. The cytokine encoded by this gene may play a role in normal lymphocyte recirculation and homing. It also plays an important role in trafficking of T cells in thymus, and in T cell and B cell migration to secondary lymphoid organs. It specifically binds to chemokine receptor CCR7.[6]

Chemokine (C-C motif) ligand 19 (CCL19) is a small cytokine belonging to the CC chemokine family that is also known as EBI1 ligand chemokine (ELC) and macrophage inflammatory protein-3-beta (MIP-3-beta). CCL19 is expressed abundantly in thymus and lymph nodes, with moderate levels in trachea and colon and low levels in stomach, small intestine, lung, kidney and spleen.[7] The gene for CCL19 is located on human chromosome 9.[8] This chemokine elicits its effects on its target cells by binding to the chemokine receptor chemokine receptor CCR7.[7] It attracts certain cells of the immune system, including dendritic cells and antigen-engaged B cells,[9][10] CCR7+ central-memory T-Cells.[11]" ([4]; `en.wikipedia.org/wiki/CCL19`)

**Freehand:** "(C-C motif chemokine 19) Chemokine secreted by dendritic cells and stromal cells in the T cell zones of lymph nodes. Attract T cells by binding to chemokine receptor 7 (CCR7)"

### 2 Supplementary Tables

S1 AAS–DESC classification performance across CLASP and baseline models . . . . . 6
S2 Ranking performance of CLASP across different description types for AAS retrieval 7
S3 Performance of different structure embeddings for clustering proteins by family . . . 8
S4 Per-family cluster distribution metrics for CLASP structure embeddings . . . . . . 9
S5 Number of AAS entries associated with each protein family . . . . . . . . . . . . . . 10
S6 Performance of different sequence embeddings for clustering proteins by family. . . 11
S7 Effect of loss-weighting strategies across alignment tasks . . . . . . . . . . . . . . . . 12
S8 Aggregate ranking of loss-weighting strategies by MCC . . . . . . . . . . . . . . . . 13
S9 Effect of removing protein names on description–structure alignment . . . . . . . . 14
S10 Effect of removing protein names on sequence–description alignment . . . . . . . . . 15
S11 Effect of removing protein names on sequence retrieval performance . . . . . . . . . 16
S12 Hypergeometric enrichment analysis of CLASP-only successes in the AAS–PDB task 17
S13 Hypergeometric enrichment analysis of CLASP-only successes in the DESC–PDB task . . . . . . . . . . . . . . . . . . . . . . . . . . . . . . . . . . . . . . . 18

**Supplementary Table S1:** AAS–DESC classification performance across CLASP and baseline models.

| Model | Accuracy | F1 Score | AUROC | AUPRC | MCC |
|---|---|---|---|---|---|
| CLASP | **0.931 ± 0.001** | **0.932 ± 0.002** | **0.979 ± 0.001** | **0.977 ± 0.002** | **0.863 ± 0.003** |
| ProteinCLIP | 0.917 ± 0.005 | 0.916 ± 0.005 | 0.969 ± 0.001 | 0.973 ± 0.001 | 0.834 ± 0.011 |
| CLIP | 0.915 ± 0.003 | 0.918 ± 0.003 | 0.968 ± 0.002 | 0.959 ± 0.003 | 0.832 ± 0.007 |
| ProtST | 0.900 ± 0.003 | 0.899 ± 0.004 | 0.962 ± 0.002 | 0.966 ± 0.002 | 0.800 ± 0.005 |
| RF | 0.829 ± 0.006 | 0.821 ± 0.007 | 0.919 ± 0.005 | 0.908 ± 0.007 | 0.661 ± 0.013 |
| LR | 0.788 ± 0.007 | 0.793 ± 0.008 | 0.867 ± 0.004 | 0.839 ± 0.002 | 0.576 ± 0.014 |

**Note:** Values are reported as mean ± standard deviation across three random seeds. AUROC = Area Under the Receiver Operating Characteristic Curve, AUPRC = Area Under the Precision–Recall Curve, MCC = Matthews Correlation Coefficient, RF = Random Forest, LR = Logistic Regression. Bold indicates the best values for each column.

**Supplementary Table S2:** Ranking performance of CLASP across different description types for AAS retrieval.

| **Protein** | **UniProt description** | **Literature description** | **Freehand description** |
|---|---|---|---|
| TAP2 | 99.99% | 99.56% | 97.62% |
| COX2 | 99.42% | 98.86% | 97.99% |
| MMP9 | 100.00% | 99.91% | 100.00% |
| CCL19 | 99.99% | 99.99% | 100.00% |
| Mean | 99.85% ± 0.25% | 99.58% ± 0.45% | 98.90% ± 1.11% |

**Note:** Mean values are reported as mean ± standard deviation across the four proteins. Per-protein percentiles indicate the position of the correct amino acid sequence in the ranked list of $n = 35,911$ candidates.

**Supplementary Table S3:** Performance of different structure embeddings for clustering proteins by family.

| **Model** | **Sil.** | **DB** | **CH** | **KL** | **JS** |
|---|---|---|---|---|---|
| CLASP | **0.15** | **2.33** | **897.73** | **2.99** | **0.54** |
| Progres | 0.08 | 3.57 | 316.15 | 2.63 | 0.43 |
| ProstT5 | 0.06 | 2.95 | 109.12 | 2.60 | 0.50 |
| COLLAPSE | -0.05 | 4.88 | 20.52 | 0.23 | 0.22 |

**Note:** Values represent clustering quality across models using centroid-based metrics. Sil. = Silhouette score; DB = Davies–Bouldin index; CH = Calinski–Harabasz index. KL = Kullback–Leibler divergence; JS = Jensen–Shannon divergence; computed between the inter- and intra-cluster distributions. For all metrics, except DB, a higher value is better. Bold indicates the best value in each column.

**Supplementary Table S4:** Per-family cluster distribution metrics for CLASP structure embeddings.

| Cluster | Intra-dist | Inter-dist | KL | JS |
|---|---|---|---|---|
| Kinases | 0.410 ± 0.295 | 0.840 ± 0.212 | 4.340 | 0.548 |
| Ion Channels | 0.355 ± 0.134 | 0.770 ± 0.222 | 2.935 | 0.674 |
| GPCRs | 0.432 ± 0.175 | 0.869 ± 0.192 | 2.516 | 0.623 |
| HSPs | 0.241 ± 0.221 | 0.949 ± 0.254 | 9.183 | 0.692 |
| ABC Transporters | 0.412 ± 0.161 | 0.779 ± 0.221 | 3.173 | 0.556 |

**Note:** Values represent clustering quality across clusters using centroid-based metrics. Intra-dist values represent the mean ± standard deviation of cosine distances from each point in a cluster to its own centroid. Inter-dist values represent the mean ± standard deviation of cosine distances from points in other clusters to that same centroid. KL = Kullback–Leibler divergence; JS = Jensen–Shannon divergence; computed between the inter- and intra-cluster distributions. All p-values from a two-sided Mann–Whitney U tests comparing intra- and inter-cluster distances were statistically significant.

**Supplementary Table S5:** Number of AAS entries associated with each protein family.

| Protein Family | Number of AASs |
|---|---|
| Kinases | 1038 |
| Ion Channels | 67 |
| GPCRs | 283 |
| HSPs | 53 |
| ABC Transporters | 49 |

**Note:** Reported counts indicate the number of unique amino acid sequences (AASs) associated with each protein family used in clustering analyses.

**Supplementary Table S6:** Performance of different sequence embeddings for clustering proteins by family.

| Model | Sil. | DB | CH | KL | JS |
|---|---|---|---|---|---|
| CLASP embeddings | **0.107** | **1.962** | **80.394** | **5.497** | **0.558** |
| ProtT5 embeddings | 0.009 | 2.732 | 49.008 | 3.410 | 0.448 |

**Note:** Values represent clustering quality across models using centroid-based metrics. Sil. = Silhouette score; DB = Davies–Bouldin index; CH = Calinski–Harabasz index. KL = Kullback–Leibler divergence; JS = Jensen–Shannon divergence; computed between the inter- and intra-cluster distributions. For all metrics, except DB, a higher value is better. Bold indicates the best value in each column.

**Supplementary Table S7:** Effect of loss-weighting strategies on CLASP performance across alignment tasks. Weights are reported in the order AAS–PDB; DESC–PDB; AAS–DESC.

| Task | Weights | Accuracy | F1 Score | AUROC | AUPRC | MCC |
|---|---|---|---|---|---|---|
| AAS–PDB | Balanced (CLASP) | 0.9160 | 0.9166 | 0.9724 | 0.9741 | 0.8321 |
| | 0 – 1/2 – 1/2 | 0.8409 | 0.8504 | 0.9205 | 0.9134 | 0.6874 |
| | 1/2 – 0 – 1/2 | 0.9130 | 0.9136 | 0.9712 | 0.9728 | 0.8261 |
| | 1/2 – 1/2 – 0 | 0.9133 | 0.9144 | 0.9711 | 0.9727 | 0.8269 |
| | 1/2 – 1/4 – 1/4 | 0.7170 | 0.7541 | 0.8207 | 0.8161 | 0.4554 |
| | **1/4 – 1/2 – 1/4** | **0.9210** | **0.9207** | **0.9731** | **0.9741** | **0.8420** |
| | 1/4 – 1/4 – 1/2 | 0.7178 | 0.7532 | 0.8199 | 0.8159 | 0.4548 |
| | 1 – 0 – 0 | 0.7328 | 0.7626 | 0.8315 | 0.8280 | 0.4812 |
| | 0 – 1 – 0 | 0.8373 | 0.8495 | 0.9215 | 0.9142 | 0.6837 |
| | 0 – 0 – 1 | 0.8840 | 0.8878 | 0.9572 | 0.9563 | 0.7697 |
| DESC–PDB | **Balanced (CLASP)** | **0.7586** | **0.7854** | **0.8519** | **0.8399** | **0.5343** |
| | 0 – 1/2 – 1/2 | 0.6303 | 0.7054 | 0.7194 | 0.6934 | 0.3032 |
| | 1/2 – 0 – 1/2 | 0.6762 | 0.7253 | 0.7712 | 0.7654 | 0.3775 |
| | 1/2 – 1/2 – 0 | 0.6872 | 0.7275 | 0.7738 | 0.7677 | 0.3921 |
| | 1/2 – 1/4 – 1/4 | 0.7396 | 0.7684 | 0.8288 | 0.8134 | 0.4949 |
| | 1/4 – 1/2 – 1/4 | 0.4998 | 0.6657 | 0.4948 | 0.4874 | -0.0000 |
| | 1/4 – 1/4 – 1/2 | 0.7368 | 0.7677 | 0.8285 | 0.8128 | 0.4914 |
| | 1 – 0 – 0 | 0.7396 | 0.7708 | 0.8376 | 0.8228 | 0.4982 |
| | 0 – 1 – 0 | 0.6218 | 0.7073 | 0.7219 | 0.6984 | 0.3004 |
| | 0 – 0 – 1 | 0.6830 | 0.7299 | 0.7736 | 0.7646 | 0.3905 |
| AAS–DESC | Balanced (CLASP) | 0.9311 | 0.9319 | 0.9777 | 0.9746 | 0.8625 |
| | 0 – 1/2 – 1/2 | 0.9322 | 0.9314 | 0.9804 | 0.9792 | 0.8648 |
| | 1/2 – 0 – 1/2 | 0.9344 | 0.9349 | 0.9795 | 0.9777 | 0.8690 |
| | 1/2 – 1/2 – 0 | 0.9342 | 0.9348 | 0.9795 | 0.9778 | 0.8685 |
| | 1/2 – 1/4 – 1/4 | 0.8018 | 0.8142 | 0.8902 | 0.8867 | 0.6091 |
| | 1/4 – 1/2 – 1/4 | 0.4993 | 0.6660 | 0.4895 | 0.4866 | -0.0222 |
| | 1/4 – 1/4 – 1/2 | 0.8021 | 0.8148 | 0.8901 | 0.8867 | 0.6099 |
| | 1 – 0 – 0 | 0.8004 | 0.8135 | 0.8902 | 0.8870 | 0.6069 |
| | **0 – 1 – 0** | **0.9350** | **0.9354** | **0.9803** | **0.9796** | **0.8701** |
| | 0 – 0 – 1 | 0.9339 | 0.9339 | 0.9796 | 0.9785 | 0.8678 |

**Note:** AUROC = Area Under the Receiver Operating Characteristic Curve, AUPRC = Area Under the Precision–Recall Curve, MCC = Matthews Correlation Coefficient.

**Supplementary Table S8:** Aggregate ranking of loss-weighting strategies based on MCC across the three alignment tasks (AAS–PDB, DESC–PDB, and AAS–DESC). Weights are reported in the order AAS–PDB; DESC–PDB; AAS–DESC.

| **Weights** | **AAS–PDB** | **DESC–PDB** | **AAS–DESC** | **Mean Rank ± Std** |
|---|---|---|---|---|
| **Balanced (CLASP)** | 2 | 1 | 6 | **3.00 ± 2.16** |
| 0 – 1/2 – 1/2 | 6 | 8 | 5 | 6.33 ± 1.25 |
| 1/2 – 0 – 1/2 | 4 | 7 | 2 | 4.33 ± 2.05 |
| 1/2 – 1/2 – 0 | 3 | 5 | 3 | 3.67 ± 0.94 |
| 1/2 – 1/4 – 1/4 | 9 | 3 | 8 | 6.67 ± 2.62 |
| 1/4 – 1/2 – 1/4 | 1 | 10 | 10 | 7.00 ± 4.24 |
| 1/4 – 1/4 – 1/2 | 10 | 4 | 7 | 7.00 ± 2.45 |
| 1 – 0 – 0 | 8 | 2 | 9 | 6.33 ± 3.09 |
| 0 – 1 – 0 | 7 | 9 | 1 | 5.67 ± 3.40 |
| 0 – 0 – 1 | 5 | 6 | 4 | 5.00 ± 0.82 |

**Note:** Rankings are computed independently for each task using MCC, with rank 1 indicating the best-performing weighting. The final column reports the mean rank and standard deviation across tasks. Bold indicates the best overall mean rank.

**Supplementary Table S9:** Effect of removing protein names from natural language descriptions on description–structure (DESC–PDB) alignment performance for CLASP.

| Model | Accuracy | F1 Score | AUROC | AUPRC | MCC |
|---|---|---|---|---|---|
| CLASP (w names) | 0.770 ± 0.011 | 0.791 ± 0.005 | 0.858 ± 0.007 | 0.846 ± 0.007 | 0.552 ± 0.018 |
| CLASP (no names) | 0.729 ± 0.034 | 0.762 ± 0.020 | 0.820 ± 0.021 | 0.809 ± 0.017 | 0.475 ± 0.059 |

**Note:** Values are reported as mean ± standard deviation across three random seeds. AUROC = Area Under the Receiver Operating Characteristic Curve, AUPRC = Area Under the Precision–Recall Curve, MCC = Matthews Correlation Coefficient.

**Supplementary Table S10:** Effect of removing protein names from natural language descriptions on sequence–description (AAS–DESC) alignment performance for CLASP.

| Model | Accuracy | F1 Score | AUROC | AUPRC | MCC |
|---|---|---|---|---|---|
| CLASP (w names) | 0.931 ± 0.001 | 0.932 ± 0.002 | 0.979 ± 0.001 | 0.977 ± 0.002 | 0.863 ± 0.003 |
| CLASP (no names) | 0.910 ± 0.008 | 0.911 ± 0.007 | 0.970 ± 0.003 | 0.970 ± 0.003 | 0.820 ± 0.015 |

**Note:** Values are reported as mean ± standard deviation across three random seeds. AUROC = Area Under the Receiver Operating Characteristic Curve, AUPRC = Area Under the Precision–Recall Curve, MCC = Matthews Correlation Coefficient.

**Supplementary Table S11:** Effect of removing protein names from natural language descriptions on sequence retrieval performance for CLASP across different description styles.

| Model | UniProt description | Literature description | Freehand description |
|---|---|---|---|
| CLASP (w names) | 99.85% ± 0.25% | 99.58% ± 0.45% | 98.90% ± 1.11% |
| CLASP (no names) | 99.77% ± 0.25% | 99.18% ± 1.22% | 98.33% ± 2.21% |

**Note:** Mean values are reported as mean ± standard deviation across the four proteins (TAP2, COX2, MMP9, and CCL19). Per-protein percentiles indicate the position of the correct amino acid sequence in the ranked list of $n = 35,911$ candidates.

**Supplementary Table S12** Hypergeometric enrichment analysis of proteins correctly classified by CLASP but not CLIP in the AAS–PDB task.

**Note:** The table is provided as a separate .xlsx spreadsheet file. For each UniProt metadata category (including organism, keywords, and functional annotations), enrichment was tested independently using a one-sided hypergeometric test, with the background population defined as the full test set for the corresponding task. Raw p-values were corrected for multiple hypothesis testing using the Benjamini–Hochberg false discovery rate (FDR) procedure.

**Supplementary Table S13** Hypergeometric enrichment analysis of proteins correctly classified by CLASP but not CLIP in the DESC–PDB task.

**Note:** The table is provided as a separate .xlsx spreadsheet file. For each UniProt metadata category (including organism, keywords, and functional annotations), enrichment was tested independently using a one-sided hypergeometric test, with the background population defined as the full test set for the corresponding task. Raw p-values were corrected for multiple hypothesis testing using the Benjamini–Hochberg false discovery rate (FDR) procedure.

### 3 Supplementary Figures

S1 Sequence-structure histograms of similarity-score distributions on the held-out test set for each baseline model . . . . . 20
S2 Description-structure histograms of similarity-score distributions on the held-out test set for each baseline model . . . . . 21
S3 Temperature ablation of CLASP alignment performance across modalities . . . . . 22

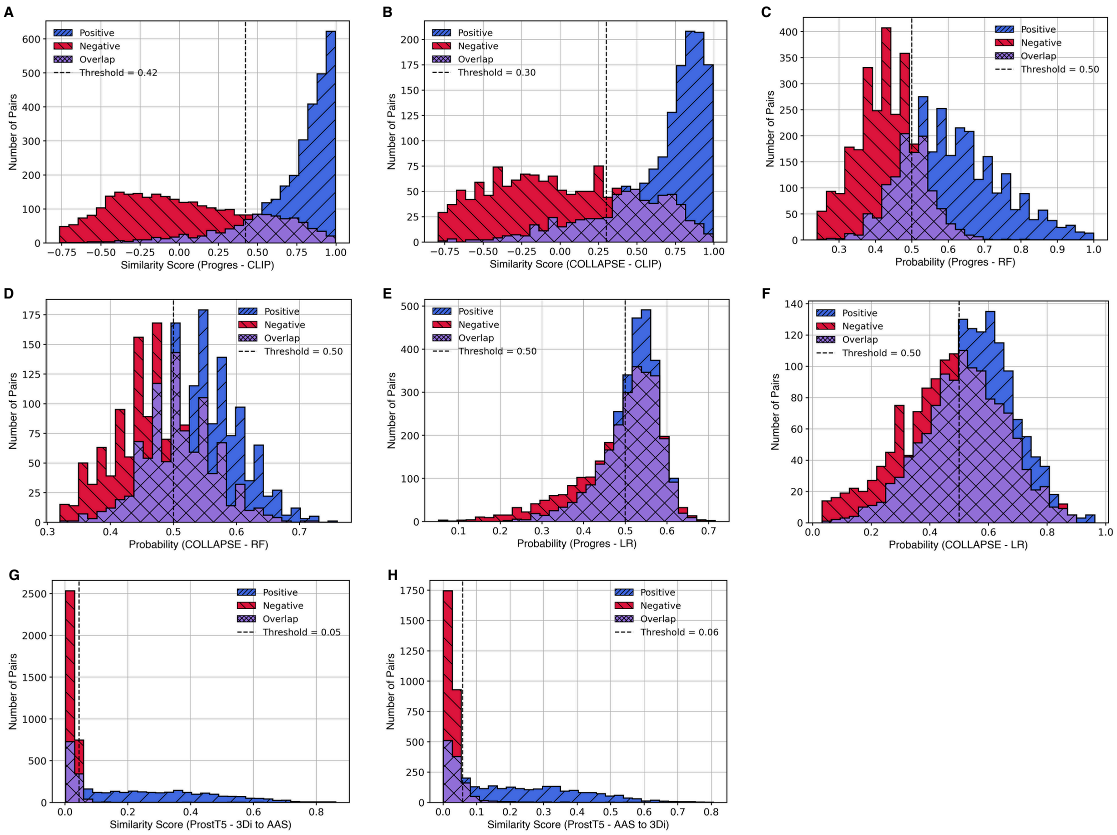

**Supplementary Figure S1:** Sequence-structure histograms of similarity-score distributions on the held-out test set for each baseline model. Panels show: Progres-CLIP **(A)**, COLLAPSE-CLIP **(B)**, Progres-RF **(C)**, COLLAPSE-RF **(D)**, Progres-LR **(E)**, COLLAPSE-LR **(F)**, ProstT5 - 3Di to AAS **(G)**, and ProstT5 - AAS to 3Di **(H)**. In each plot, positive pairs are shown in blue with diagonal hatching, negative pairs in red with vertical hatching, and overlapping regions in purple with cross-hatching. The black dashed line indicates the optimal decision threshold calculated using the validation set.

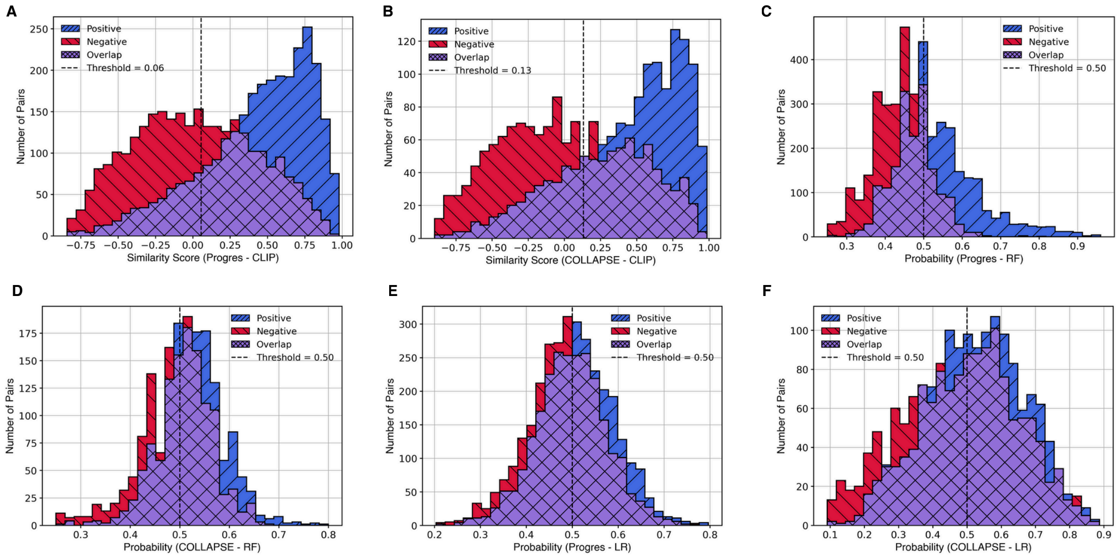

**Supplementary Figure S2:** Description-structure histograms of similarity-score distributions on the held-out test set for each baseline model. Panels show: Progres-CLIP **(A)**, COLLAPSE-CLIP **(B)**, Progres-RF **(C)**, COLLAPSE-RF **(D)**, Progres-LR **(E)**, and COLLAPSE-LR **(F)**. In each plot, positive pairs are shown in blue with diagonal hatching, negative pairs in red with vertical hatching, and overlapping regions in purple with cross-hatching. The black dashed line indicates the optimal decision threshold calculated using the validation set.

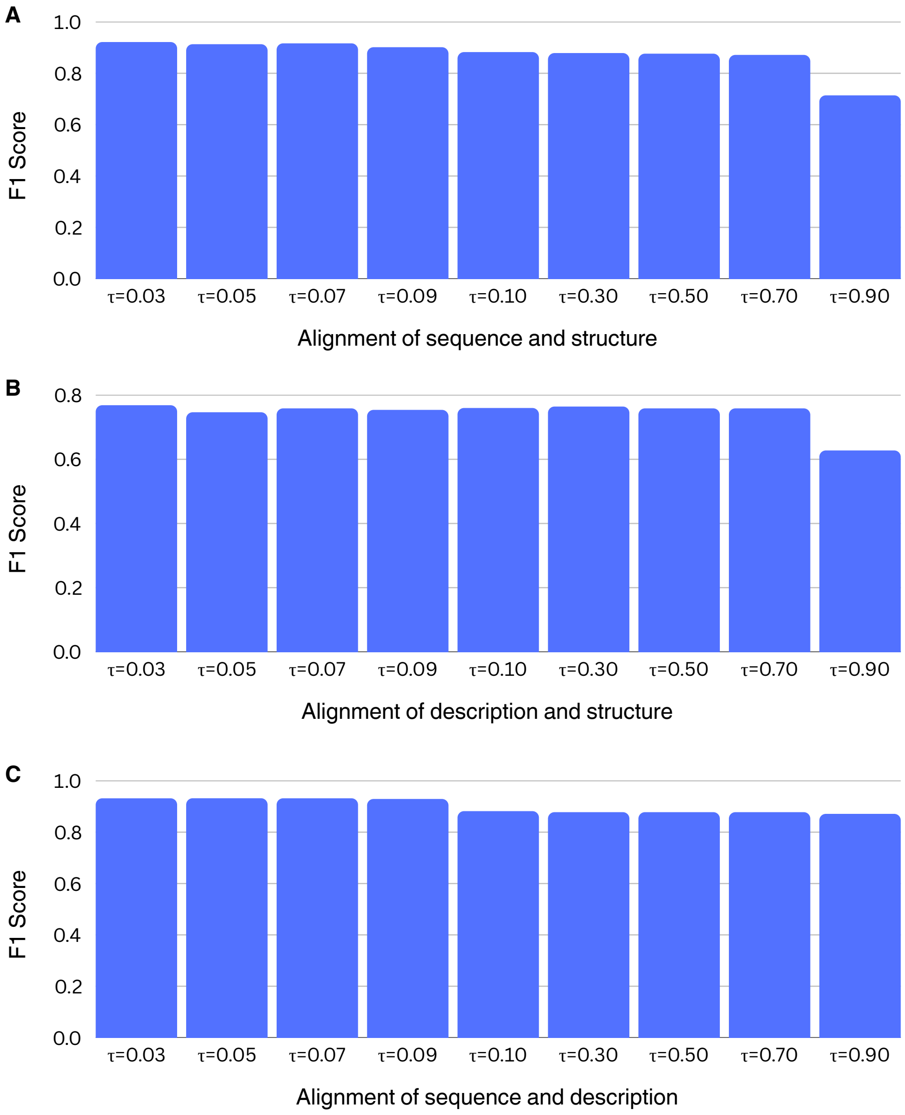

**Supplementary Figure S3:** Effect of the contrastive temperature parameter $\tau$ on CLASP alignment performance across modalities, measured by F1 score on the held-out test set. **(A)** Alignment of sequence and structure (AAS–PDB), **(B)** alignment of description and structure (DESC–PDB), and **(C)** alignment of sequence and description (AAS–DESC).
